# Protective effects of a novel RIPK3 antagonist against neutrophil necroptosis and formation of neutrophil extracellular traps

**DOI:** 10.64898/2026.07.20.739512

**Authors:** Eugen Koren, Erik Foehr, Tatjana Faruqi, Elisabeth Gardiner, Rajiv Mahadevan

## Abstract

Functionally competent, highly purified neutrophils were isolated from healthy human donors. Neutrophil necroptosis and neutrophil extracellular trap (NET) formation were induced using TNF-α in combination with the pan-caspase inhibitor zVAD-FMK and the IAP antagonist BV6. NETs were visualized, quantified, and morphologically characterized by fluorescence microscopy following staining with Hoechst 33342 and Sytox Green. Levels of extracellular, cell-free neutrophil elastase (NE), myeloperoxidase (MPO), and DNA were also measured as indicators of NET release. The ability of TACT507, a proprietary RIPK3 antagonist, to efficiently block NETs formation and NET related necroptosis was evaluated. TACT507 demonstrated concentration dependent, significant inhibition of neutrophil necroptosis and NETs formation.

**Summary Sentence:** Isolated human neutrophils treated *ex vivo* by TNF-α in combination with apoptosis inhibitors, underwent necroptosis causing formation of neutrophil extracellular traps. This process was inhibited by the proprietary antagonist of RIPK3.

## Introduction

Neutrophil extracellular traps (NETs) are web-like structures composed of DNA, histones, and granule proteins extruded from neutrophils. There are two mechanisms of NETs formation, NETosis and necroptosis, both ultimately leading to regulated cell death. NETosis is primarily driven by the ROS-PAD4 axis (1) and is uniquely hardwired into neutrophils as a defense mechanism to actively trap and kill pathogens. Necroptosis is driven by the RIPK1/RIPK3/MLKL pathway to eliminate compromised cells of any type including neutrophils, epithelial, endothelial, and parenchymal tissue cells. While NETosis is optimally triggered by contact with live microbes or pathogen-associated molecular patterns (PAMPs) like LPS (2), necroptosis is induced by endogenous, host related non-infectious factors such as cytokines and various ligands (e.g. TNF-α, Interferons, FasL, TRAIL) and by physiological stressors (reactive oxygen species, ATP depletion (3-5). The signal transduction cascade resulting in necroptotic cell death is initiated by recruitment of the Receptor-Interacting Protein Kinase 1, (RIPK1), which phosphorylates Receptor-Interacting Protein Kinase 3 (RIPK3). Further signaling steps include phosphorylation of mixed lineage kinase domain-like (MLKL) protein by RIPK3, resulting in its oligomerization, translocation to the inner leaflet of the plasma membrane, causing a membrane rupture and formation of NETs (3).

Given the abundance of neutrophils and their ability to rapidly recruit to sites of tissue injury, neutrophils play a substantial role in inflammatory and degenerative disease states (6). Necroptotic NETs can either trigger or significantly augment existing necroptosis by releasing tissue damaging proteolytic enzymes, histones and free radicals as well as damage-associated molecular patterns (DAMPs) such as high-mobility group box 1 (HMGB1), S100 proteins and extracellular DNA (7). DAMPs bind to pattern recognition receptors on nearby healthy cells as well as newly recruited neutrophils enhancing further necroptosis. This continuous cycle of necroptotic cell death perpetuates inflammation and tissue injury with neutrophils playing a prominent exacerbating role.

In this study we demonstrate protective effects against neutrophil necroptosis and NETs formation achieved by TACT507, a proprietary compound designed as an antagonist of RIPK3, a key factor in the necroptotic signaling cascade.

## Methods

### Isolation of neutrophils

Highly purified and viable neutrophils were isolated from two healthy human blood donors (one female and one male) screened and tested according to the Association for the Advancement of Blood & Biotherapies (AABB) guidelines and FDA requirements at BioIVT lab in Berkeley, California. Neutrophil isolation was approved by BioIVT’s institutional review board (IRB). Briefly, one unit of the whole blood (250 mL) was processed by Miltenyi’s magnetic bead negative selection method. The resulting neutrophil suspension was centrifuged at 400 rpm and 4-8°C, washed and re-suspended in 50 mL of cold (4-8°C) 1640 RPMI with no phenol red, spiked with penicillin/streptomycin and supplemented with 10% mycoplasma free FBS. This was followed by flow cytometry using a sequential gating strategy including the side scatter/forward scatter, side scatter/CD45 and CD 16/66b gates to assess neutrophil purity. The trypan blue exclusion test was carried out to determine cell viability. The final cell number was adjusted to 150×10^3^ per mL.

### Reagents

Whole Blood Neutrophil Isolation Kit, human, Miltenyi Biotech Cat No 130-104-434; zVAD Pan-Caspase Inhibitor, Thermo Fisher Cat No A62645; BV6 inhibitor of apoptosis proteins (IAPs), InvivoGen Cat No inh-bv6; Recombinant human TNF-α, Origene Cat. No. TP750007; DMS0 Thermo fisher Cat No D12345; Penicillin/Streptomycin, Thermo Fisher Cat No 15140122; Mycoplasma free FBS, ThermoFisher Cat No A5256801; SYTOX Green Nucleic Acid Stain, Thermo Fisher Cat No S7020; Hoechst 33342, Fisher Cat No 62249; 1640 RPMI without phenol red, Thermo Fisher Cat No 11835030; TACT507 proprietary compound developed at Tactile Therapeutics, Inc.

### Neutrophil incubation procedures

Two separate incubation experiments were conducted using neutrophils from both donors, one to detect NETs after staining with Hoechst 33324 and one to detect NETs after staining with Sytox Green. For staining with Hoechst, neutrophils were seeded in a 24-well plate (AG Advangene, CC Plate-PS-96S-F-C-S) with 50,000 cells per well and incubated for 18 hours in a humidified CO2 incubator at 37°C. Six wells were used to treat neutrophils as follows: A) with incubation in medium alone; B) incubation medium + TACT507 compound at 3 μM concentration; C) incubation medium + necroptotic cocktail alone containing 25 μM zVad, 5 μM BV6 in 0. 4% DMSO and 100 ng TNF-*α* as final concentrations; D) incubation medium + necroptotic cocktail and TACT507 at 3 μM concentration; E) incubation medium + necroptotic cocktail and TACT507 at 1 μM concentration; F) incubation medium + necroptotic cocktail and TACT507 at 0.3 μM concentration. Wells spiked with zVAD, and TACT507 were pre-incubated at RT for 1 hour to allow for their uptake by neutrophils. After the pre-incubation, BV6 and TNF-*α* were added at the same time. Immediately after incubation, 100 μL of Hoechst 33342 diluted 1:2000 in PBS was directly added to each well without fixation. After 15 minutes of staining at room temperature, all wells were scanned with Invitrogen’s FLoid Cell Imaging Station (Catalog No. 10213412) using the blue fluorescence emission channel.

To quantify protective effects of TACT507 against neutrophil necroptosis and NET formation, microscopic and biochemical analyses were carried out after the 18-hour incubation of neutrophils in a humidified CO2 incubator at 37°C. As described above, wells spiked with zVAD, and TACT507 were pre-incubated at RT for 1 hour to allow for their uptake by neutrophils. After the pre-incubation, BV6 and TNF-*α* were added at the same time. A combination of Sytox Green staining and plate-based assays for the released DNA, neutrophil elastase (NE) and myeloperoxidase (MPO) was carried out. Neutrophils were incubated in 96-well plates (AG Advangene, CC Plate-PS-96S-F-C-S). Incubation mixtures were prepared using duplicate wells, each containing 200 μL neutrophil suspension with 30,000 cells per well. After the incubation, supernatants from each well were carefully collected avoiding cells at the bottom and pooled for the cell-free DNA, NE and MPO analyses after centrifugation at 1500 rpm for 10 minutes. 100 uL of the PBS buffer containing 4% paraformaldehyde was added to each well to fix cells and prevent drying of attached cells and NETs. Each well was then stained with 100 μL of 1 μM Sytox Green for 15 minutes protected from light and scanned with Invitrogen’s Floid Cell Imaging Station using the green fluorescence emission channel. All incubation mixtures contained 0.4% DMSO and were handled under sterile conditions.

### Microscopic analyses

After the staining with Sytox Green, each well was randomly scanned by two independent analysts using Invitrogen’s Evos FLoid Imager under the green fluorescence channel. Multiple scans were taken from all wells to determine the number of non-lysed, round-shaped neutrophils using the Image J software. NETs were enumerated by visual counting carried out by two independent analysts achieving a positive correlation between their respective NET counts (Pearson correlation coefficient r=0.76). Numbers of non-lysed neutrophils and NETs were pooled from all available scans to calculate ratios between NETs and non-lysed neutrophils (total number of NETs divided by total number of non-lysed neutrophils). All incubation mixtures contained 0.4% DMSO to ensure solubility of zVAD, BV6, and TACT507. Incubations were carried out under sterile conditions.

### Biochemical analyses

To measure cell-free DNA released from neutrophils, we used the SYTOX Green induced fluorescence measured with the Molecular Devices iD3 plate reader. SYTOX® Green (Invitrogen Cat. No S7020, 5 nM in DMSO), is a high-affinity nucleic acid stain that induces a >500-fold fluorescence enhancement upon binding to DNA. A dsDNA plasmid (pGMPMax, Lonza) was used as a standard to generate a calibration curve based on the 10000, 5000, 2500, 1250, 625, 312.5, and 156.25 ng/mL concentration range. Test samples representing supernatants from the neutrophil incubation mixtures were transferred to the black-walled 96 well plates (Olympus Plastics, Cat#22-721). Diluted (1:1000) SYTOX Green was added to the standards and test samples and incubated for 10 minutes protected from light. This was followed by the fluorescence measurement at the 488 nm excitation and 523 nm emission wavelength. DNA concentrations were extrapolated from the calibration curve.

Neutrophil elastase and myeloperoxidase released into neutrophil culture supernatants were quantified with commercial assays for NE (Abcam Cat No ab 204730) and MPO (Invitrogen Cat No BMS2038INST).

## Results

### Isolation of neutrophils

Isolation of human primary neutrophils from whole blood drawn from two healthy blood donors resulted in highly purified and viable cells as shown in Figures 1 and 2. A compact population of cells was found in a typical side/forward scatter position as well as the side scatter/CD45 and the CD16/CD66b gates. This gating strategy revealed a homogenous population of highly purified (98%) mature neutrophils (Figure 1). In the trypan blue exclusion test, most cells were translucent and clean of trypan blue demonstrating high degree of viability (Figure 2). High level purity and viability of isolated neutrophils was virtually identical in both donors.

**Figure 1.**
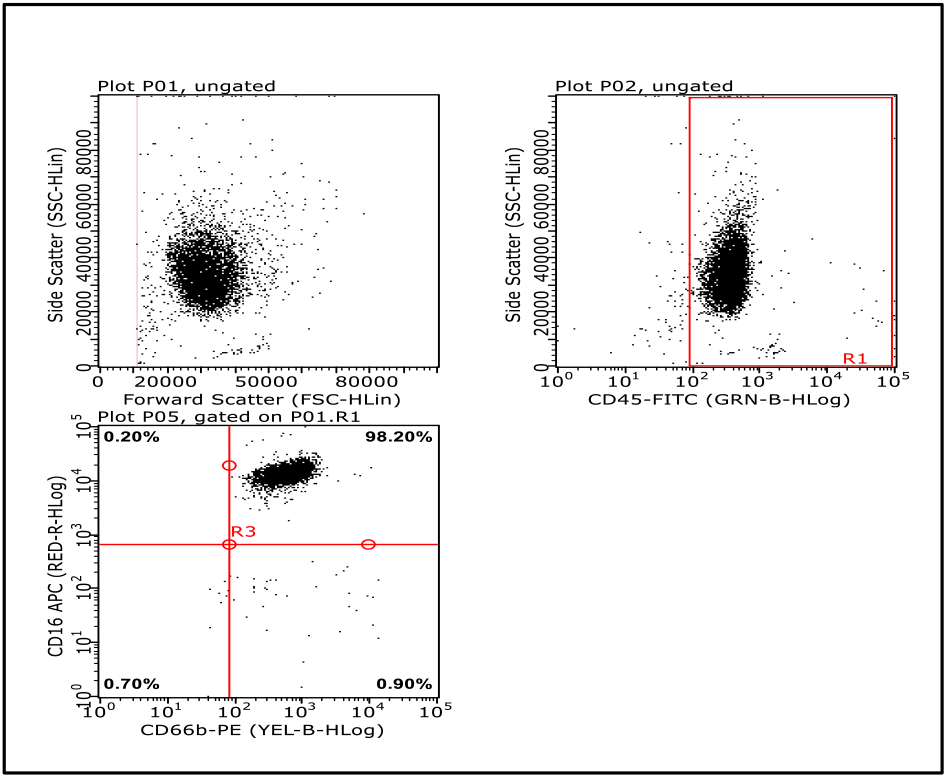
Flow ctyomerty analysis of neutrophils isolated from the whole blood of a healthy donor. Compact populations of cells were found in a typical side/forward scatter position as well as in the side scatter/CD45 and the CD16/CD66b gates. This gating strategy revealed a homogenous population of highly purified (98%) mature neutrophils. This pattern was virtually identical in both blood donors.

**Figure 2.**
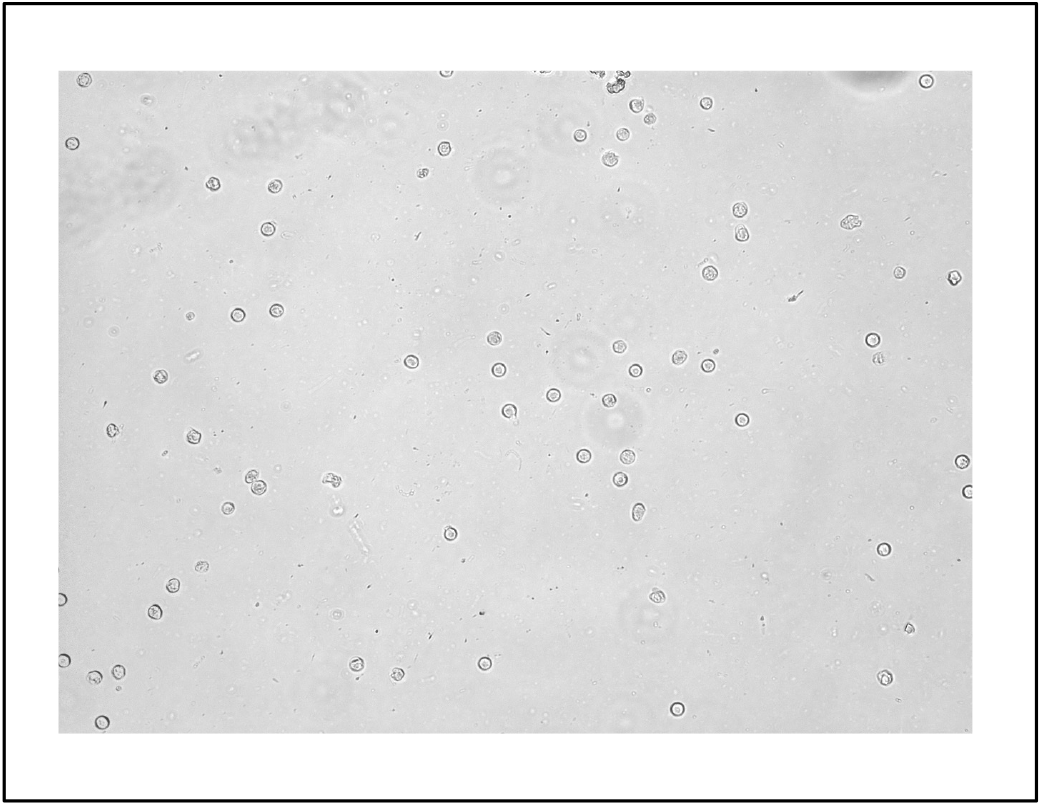
Viability of isolated neutrophils. Representative bright field image of freshly isolated primary neutrophils after trypan blue exclusion test showing overwhelming number of translucent, round-shaped cells without presence of the trypan blue demonstrating high degree of the cell viability.

### Effects of the necroptotic cocktail (zVAD/BV6/ TNF-α) and TACT507 on NETs formation

Incubation of purified neutrophils in the cell culture medium alone showed multiple single, non-lysed cells after staining with Hoechst 33342 (Figure 3A). Single cells were also detected after incubation with 3 μM TACT507 alone, showing no visible effects on neutrophil morphology (Figure 3B). Incubation with the necroptotic cocktail alone resulted in pronounced formation of large masses stained with Hoechst representing NETs (Figure 3C). This effect was greatly diminished in the well incubated with the necroptotic cocktail and 3 μM TACT507 where single cells mixed with some smaller, but no large NETs were found (Figure 3D). In the well incubated with the necroptotic cocktail and 1 μM TACT507, a few large and some smaller NETs mixed with occasional singe cells were visible (Figure 3E). A large NET with multiple single neutrophils were found in the well incubated with the necroptotic cocktail and 0.3 μM TACT507 (Figure 3F). These patterns indicated concentration dependent protective effects of TACT507 against necroptosis-induced NETs formation. TACT507 alone showed no apparent effect on neutrophil morphology. A more detailed view of NETs after staining with Sytox Green revealed web-like structures formed by extruded DNA strings as shown in Figure 4.

**Figure 3.**
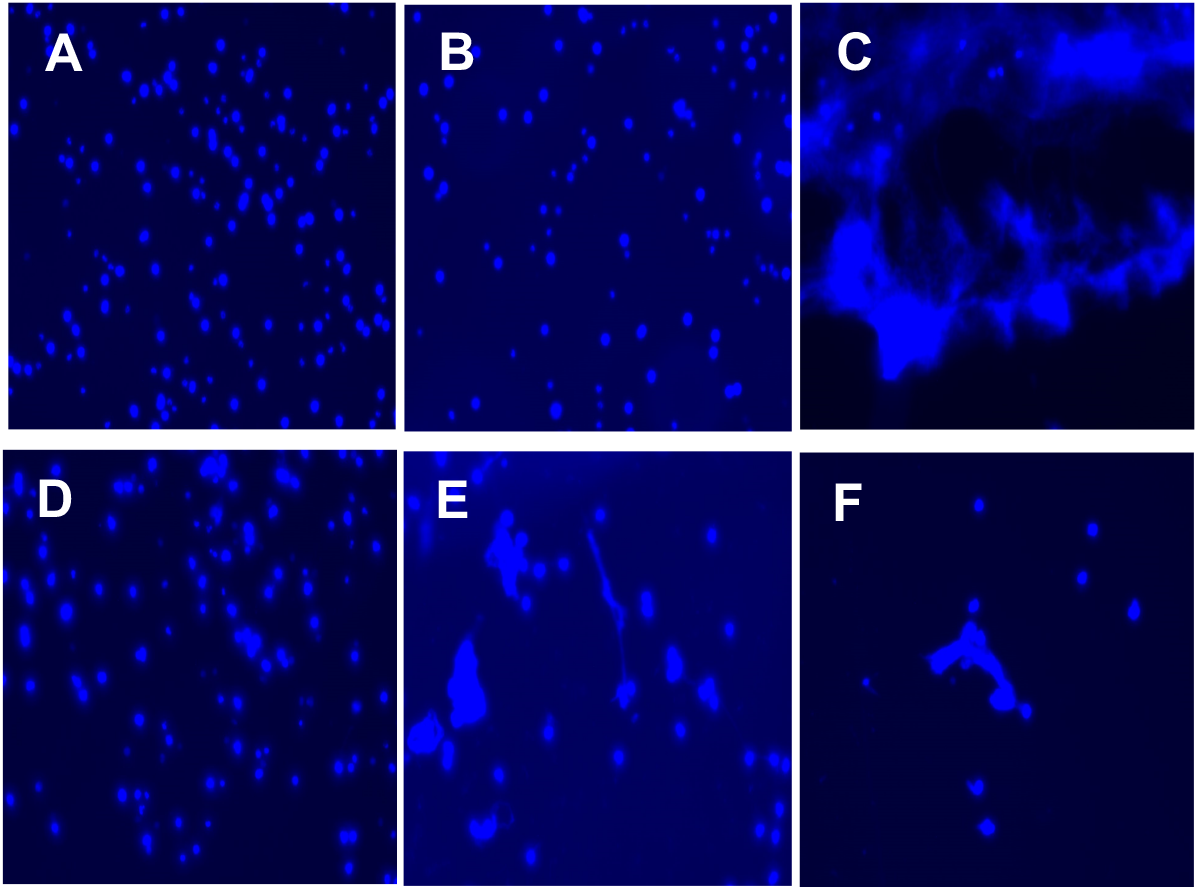
Effect of TACT507 on NET formation in response to 18-hour zVAD/BV6/TNF-α treatment. Hoechst 33342-stained neutrophils visualized by Invitrogen’s Floid Cell Imaging Station. A) No treatment. B) Treatment with 3 μM TACT507 alone. C) Treatment with zVAD/BV6/TNF-α alone. D) Treatment with zVAD/BV6/TNF-α + 3 μM TACT507. E) Treatment with zVAD/BV6/TNF-α + 1 μM TACT507. F) Treatment with zVAD/BV6/TNF-α + 0.3 μM TACT507.

**Figure 4.**
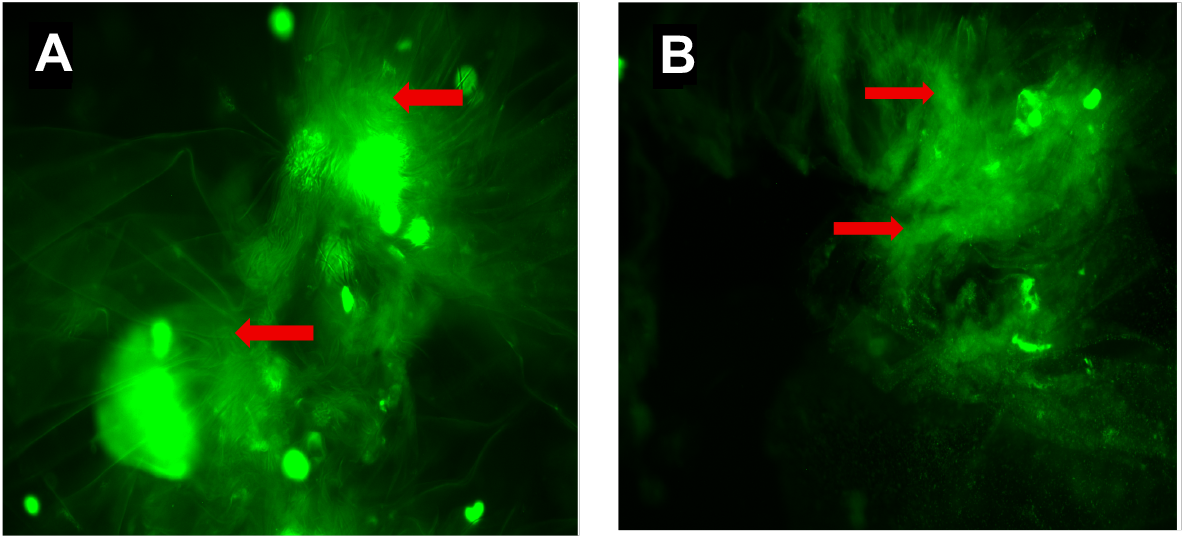
Appearance of NETs stained with Sytox Green under higher (40X) magnification. Extracellular strings of DNA (red arrows A and B), reveal web-like structures of extracellular traps.

### Quantitative assessment of the protective effect of TACT507

Microscopic analyses of neutrophils stained with Sytox Green combined with measurements of released cfDNA, NE and MPO allowed for quantitative assessment of the protective effects of TACT507 against formation of NETs. Figure 5A, represents neutrophils incubated for 18 hours in cell culture medium alone. Numerous, predominantly round shaped, non-lysed neutrophils are visible. In contrast, incubation with zVAD/BV6/TNF-α alone resulted in multiple NETs and some non-lysed neutrophils (Figure 5B). Total numbers of non-lysed neutrophils and NETs as well as ratios between NETs and non-lysed neutrophils are shown in Table 1.

**Figure 5.**
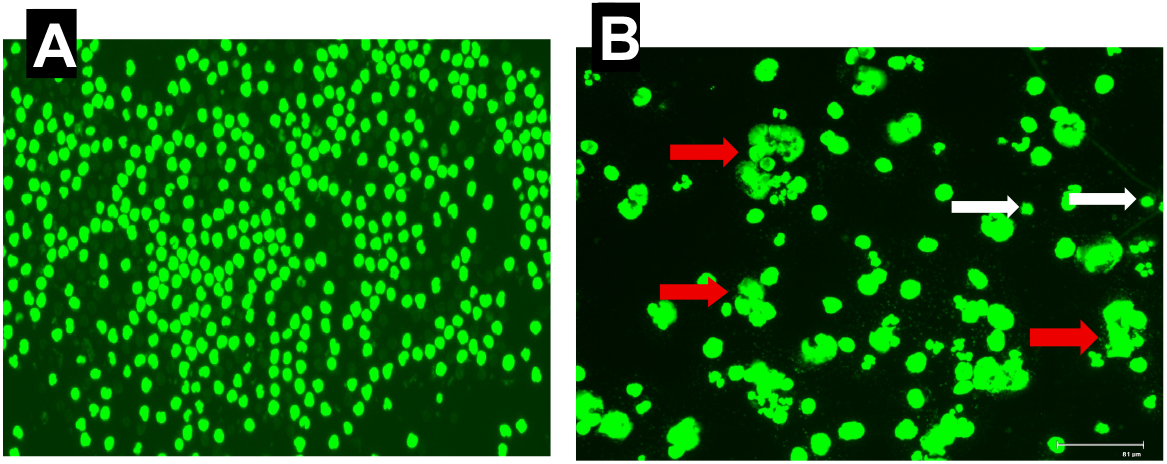
Morphology of the Sytox green-stained neutrophils and NETs. Representative images of neutrophils incubated with cell culture medium alone (A) and with zVAD/BV6/TNF-α alone (B). Numerous, predominantly round-shaped non-lysed neutrophils of uniform size are visible in panel A. Panel B shows multiple irregularly shaped large and coalescing NETs (red arrows). Rare non-lysed neutrophils (white arrows) are also seen panel B. According to the scale bar of 81 μM in panel B (bottom right), sizes of non-lysed neutrophils vary between 8 and 12 μM whereas NETs fall within the 25 to 50 μM range.

**Table 1.** Frequency and distribution of non-lysed neutrophils and NETs under various treatment conditions. Ratios between numbers of NETs and non-lysed neutrophils clearly demonstrate concentration dependent protective effects of TACT507 against necroptotic NET formation. This concentration dependent protective effect was also confirmed by significant positive correlation between numbers of non-lysed neutrophils and concentrations of TACT507 (Table 1) achieving Pearson’s correlation coefficient of r=0.979, p<0.05 (StatsMasters.com).

| Treatment Condition | Number of fields analyzed | Total number of non-lysed neutrophils | Total number NETs | Ratios between NETs and non-lysed neutrophils |  |
| --- | --- | --- | --- | --- | --- |
| Neutrophils incubated with zVAD/BV6/TNF-α alone | 7 | 6 | 28 | 4.67 | Fold of ratio decrease relative to zVAD/BV6/TNF-α alone |
| Neutrophils incubated with zVAD/BV6/TNF-α +3μM TACT507 | 21 | 146 | 47 | 0.32 | 14.59 |
| Neutrophils incubated with zVAD/BV6/TNF-α +1μM TACT507 | 20 | 58 | 30 | 0.52 | 9.98 |
| Neutrophils incubated with zVAD/BV6/TNF-α +0.3μM TACT507 | 20 | 53 | 64 | 1.21 | 3.86 |

### Measurements of released NE, MPO, and cell-free DNA

Results of microscopic analyses were supported by the measurements of released NE, MPO, and cell-free DNA. NE, MPO and cell-free DNA levels in supernatants from neutrophils treated with necroptotic cocktail alone were significantly higher in comparison with supernatants from neutrophils incubated with necroptotic cocktail and 3μM TACT507 as shown in Figure 6.

**Figure 6.**
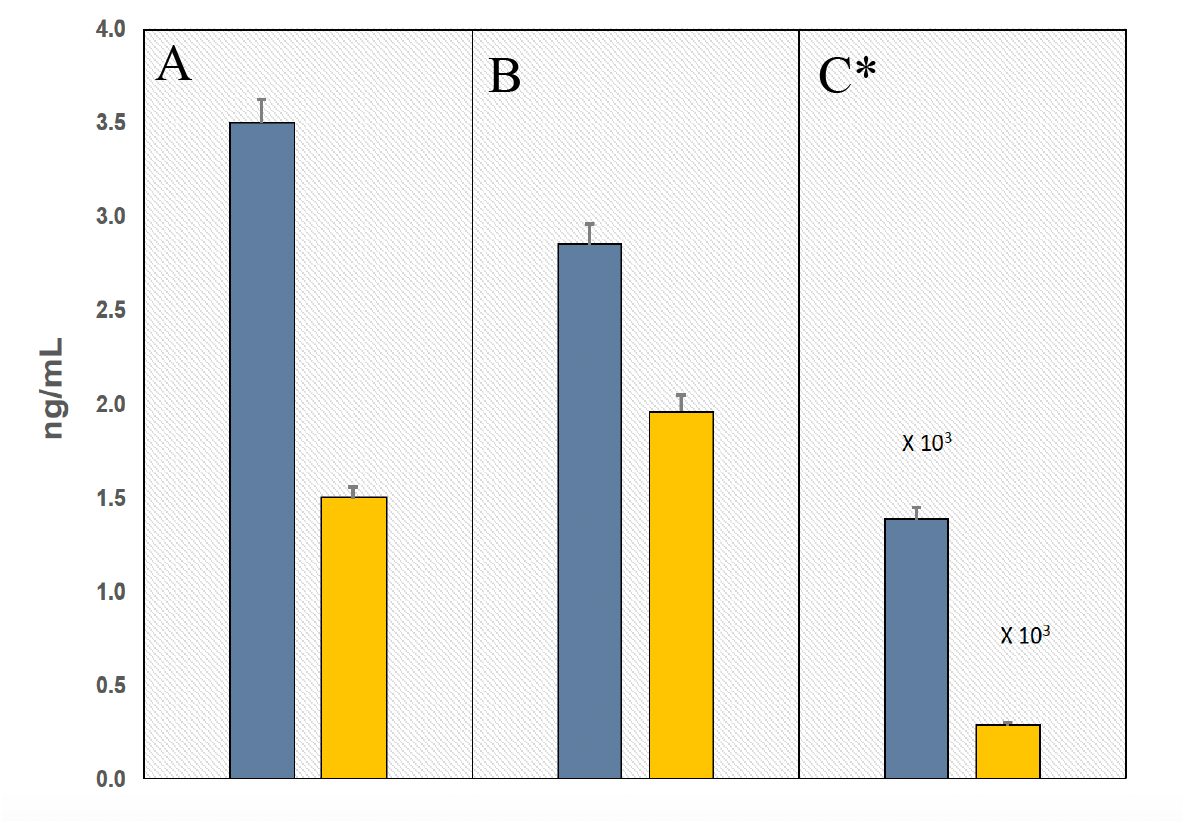
Protective effects of TACT507 against the release neutrophil elastase, myeloperoxidase and cell-free DNA. TACT507 at 3μM concentration (yellow bars) diminished the release of elastase (A); myeloperoxidase (B); and cell-free DNA (C) from neutrophils during NET formation caused by treatment with zVAD/BV6/TNF-α. Blue bars represent cell-free NE, MPO and cfDNA released from neutrophils incubated with zVAD/BV6/TNF-α alone. *cfDNA bars are shown at X 10^3^ scale. All differences were significant (p0.05) by respective paired T-tests (StatsMasters.com).

In conclusion, the zVAD/BV6/ TNF-α cocktail readily induced necroptosis and NETs formation in freshly isolated primary human neutrophils. TACT507 protected against necroptosis and NETs formation in a concentration dependent fashion at 3.0, 1.0 and 0.3 μM concentrations.

## Discussion

Human neutrophils were stimulated *ex vivo* with recombinant TNF-α in the presence of the pan-caspase inhibitor zVAD-FMK and the inhibitor-of-apoptosis (IAP) antagonist BV6. This established necroptosis-inducing cocktail effectively triggered RIPK3–MLKL–dependent neutrophil necroptosis and led to robust formation of NETs. Microscopic analysis demonstrated the characteristic lytic morphology of necroptotic NETs, including extensive extracellular DNA fibers, accompanied by elevated levels of cell-free DNA, NE and MPO in culture supernatants. These features are consistent with plasma-membrane rupture mediated by oligomerized MLKL, the terminal effector of necroptotic cell death (1,7).

Neutrophil necroptosis and necroptosis-associated NET formation have been documented in multiple human pathologies and are increasingly recognized as amplifiers of inflammation and tissue injury. Diseases such as gout (8) vasculitis (9), and inflammatory bowel disease (10) exhibit strong evidence of damaging NET involvement. Moreover, necroptosis contributes significantly to the pathogenesis of acute lung (11) and kidney injury (12), where emerging data suggests a mechanistic link between necroptotic signaling and neutrophilic NET formation.

While neutrophils typically do not cross the blood-brain barrier, they infiltrate the CNS following injury or disease, where both necroptosis and NET formation contribute to inflammation and neuronal damage (13). Increased levels of necroptosis mediators (RIPK1, RIPK3, MLKL) have been observed in post-mortem brain tissues from patients with Parkinson disease and Gaucher disease (14). NETs have been identified as mediators of cerebral edema and elevated intracranial pressure following traumatic brain injury, contributing to neurological deficits (15). They also contribute to the neurodegenerative process in Alzheimer disease and amyotrophic lateral sclerosis (ALS) (16). In cerebral stroke NETs promote thrombosis aggravating tissue damage after a stroke (17). Neutrophil extracellular traps play a detrimental role in age-related macular degeneration and retinal degeneration, where the release of NETs leads to damage to photoreceptors and retinal pigment epithelial cells, contributing to vision loss (18).

The presence of NETs in various stages of human atherosclerotic lesions (19,20) points to an important role of neutrophil necroptosis and formation of neutrophil extracellular traps the during progression of atherothrombosis, a pathological process that underlies cardiovascular diseases. NETs promote atherosclerosis by releasing harmful intracellular contents (DAMPs and proteolytic enzymes) creating a pro-inflammatory environment that activates macrophages, platelets and endothelial cells leading to plaque destabilization, thrombosis and vascular damage (21). Increased NET markers in coronary artery disease (22) carotid artery disease (23) and aortic aneurysms (24) as well as other severe complications such as peripheral artery disease (25) are linked to worse clinical outcomes in all these conditions.

Not surprisingly, numerous attempts to inhibit neutrophil necroptosis and NET formation have been pursued over the past decade. A wide range of NET-targeting strategies, including PAD4 inhibitors, DNase-based NET degradation, blockade of upstream neutrophil activation pathways, and neutralization of NET-associated cytotoxic components, have been tried, with varying success in preclinical models of inflammation and thrombotic disease (26). However, clinical translation of NET inhibitors has achieved limited success, largely due to challenges in model relevance, NET heterogeneity, off-target effects, and pharmacokinetic limitations.

In this study, we demonstrate that the proprietary TACT507 compound, designed as a novel RIPK3 antagonist, prevents *ex vivo* induced neutrophil necroptosis and NETs formation in a concentration dependent fashion. The use of human neutrophils and the translationally relevant approach of blocking the primary necroptosis mediator, RIPK3, shows encouraging results. As RIPK3 plays the central role in driving the necroptotic pathway, the use of TACT507 may serve as the basis for further development of RIPK3 antagonists aimed at protection against the neutrophil necroptosis and NETs formation as well as protection against necroptosis across a range of other cell types and tissues.

## Acknowledgments and Sources of Funding

Tactile Therapeutics, Inc. provided funding and resources for this study.

## Authorship Contribution Statement

Conceptualization: EK Investigation: EK, EF

Data Curation-Formal Analysis: EK, EF, TF

Project Administration/Oversight: TF, RM

Writing-Original Draft: EK

Writing-Review, Editing, and Revision: EK, EG, TF, EF, RM

